# A human cancer cell line initiates DNA replication normally in the absence of ORC5 and ORC2 proteins

**DOI:** 10.1101/2020.08.10.245076

**Authors:** Etsuko Shibata, Anindya Dutta

## Abstract

The Origin Recognition Complex (ORC), composed of six subunits, ORC1-6, binds to origins of replication as a ring-shaped heterohexameric ATPase that is believed to be essential to recruit and load MCM2-7 around DNA and initiate DNA replication. We had reported the creation of viable cancer cell lines that lacked detectable ORC1 or ORC2 protein without a significant decrease in the number of origins firing. We now report that human HCT116 colon cancer cells also survive with a mutation in the initiator ATG of the *ORC5* gene that abolishes the expression of ORC5 protein. Even if an internal methionine is used to produce an undetectable, N terminally deleted ORC5, the protein would lack 80% of the AAA+ ATPase domain, including the Walker A motif. The ORC5-depleted cells show normal chromatin binding of MCM2-7 and initiate replication from similar number of origins as wild type cells. In addition, we introduced a second mutation in *ORC2* in the *ORC5* mutant cells rendering both ORC5 and ORC2 proteins undetectable in the same cells, and destabilizing the ORC1, ORC3 and ORC4 proteins. Yet the double mutant cells grow, recruit MCM2-7 normally to chromatin and initiate DNA replication with normal number of origins. Thus, in these selected cancer cells, either a crippled ORC lacking ORC2 and ORC5 and present at minimal levels on the chromatin can recruit and load enough MCM2-7 to initiate DNA replication, or human cell-lines can sometimes recruit MCM2-7 to origins independent of ORC.

## Introduction

The six-subunit ORC (1) is the initiator protein that binds to replicator sequences to initiate DNA replication in eukaryotes. ORC is essential for DNA replication in yeasts, flies, mice and most likely in humans (2–8). The six-subunit, ring-shaped ORC ATPase complex co-operates with the CDC6 ATPase protein to promote the binding of CDT1 and MCM2-7 complex to DNA. MCM2-7 is a core part of the replicative helicase essential for DNA replication (1, 9–16). Unlike yeast ORC where all six subunits form a tight complex and ORC1-5 subunits have Walker A and B motifs that bind and hydrolyze ATP, the six subunits of human ORC are not associated with each other into one tight complex. ORC2-5 form a core subcomplex (17), with ORC1, the only subunit responsible for the ATPase activity of ORC (18–20), associated loosely with the core. ORC6 is very poorly associated with the core of ORC2-5 subunits. Although human ORC was expected to be essential for cell viability and proliferation, we were surprised to obtain viable clones of cells from three cell lines that were mutated on both alleles of *ORC2* or *ORC1* such that neither protein was detectably expressed (21). Although the ORC1 or ORC2 proteins were undetectable by Westerns, the surprising nature of the finding made us consider the possibility that the mutated ORC subunits were expressed at levels below the sensitivity of detection of our antibodies. Quantitative westerns, however, revealed that if these clones expressed any ORC2 protein, it was at <1500 molecules per cell, which was probably not sufficient to license the 52,000 origins of replication that were firing per cell-cycle per cell.

There were two questions that needed to be addressed. (1) Could an N terminally deleted ORC2 protein, initiated from an internal methionine, be expressed at <1500 molecules/cell and function in the loading of MCM2-7 as part of the ORC ring? (2) Could the ORC five-subunit ring be constructed by incorporating CDC6 or another ORC subunit instead of the missing ORC2 or ORC1 subunits? To address these, we have re-visited the issue by deleting the initiator methionine of *ORC5* in HCT116 colon cancer cells and selecting for viable, proliferating cells. Again, we do not see any ORC5 protein, including any short isoform, but in this case we are sure that even if an alternate isoform of the mRNA is produced to initiate translation from an internal methionine, the resulting protein would have lost most of the N terminal AAA+ ATPase lobe of ORC5, including the Walker A motif essential for ATP binding. Even more surprising, additional mutation of both alleles of *ORC2* in the *ORC5* mutant cells produced viable cell lines that showed relatively normal DNA replication even though two subunits of ORC are undetectable. The result affirms the hypothesis that either a crippled ORC missing several ORC subunits in the ATPase ring can recruit MCM2-7 to DNA or that human cancer cells can be selected that have bypassed the requirement of ORC for MCM2-7 recruitment and replication initiation.

## Results

### Biallelic disruption of the *ORC5* gene

The first methionine in the *ORC5* mRNA is located in exon 1 and is the initiator methionine for the protein. If this methionine is deleted, the next internal methionine is located at position 133, downstream from the Walker A site required to bind ATP (**Fig. 1A**). In fact if a protein is produced from methionine 133, it will have deleted most of the AAA+ ATPase lobe of ORC5, a lobe that is critical for interaction with the other AAA+ ATPase lobes in the five-subunit ORC ring (9). CRISPR-Cas9 mediated DNA break was directed at the first ATG of *ORC5* in HCT116 p53-/-cells and single cell clones screened by immunoblotting for ORC5 protein. Three clones where no ORC5 protein was detected were genotyped by amplifying the region around the ATG and sequencing (**Fig. 1B**). All three clones suffered bi-allelic mutation in the *ORC5* gene that deleted the first methionine.

**Figure 1.**
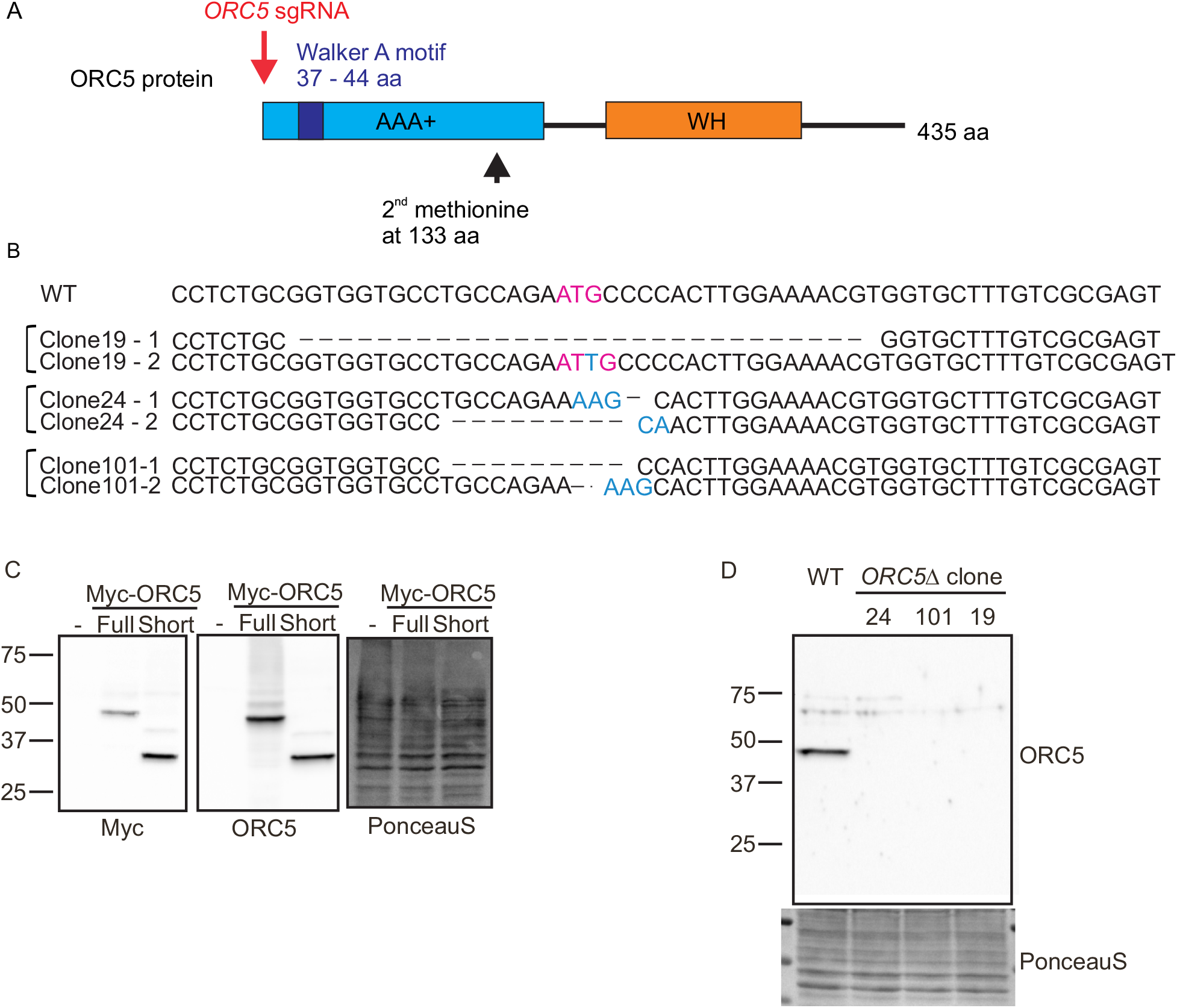
Deletion of *ORC5* in HCT116 p53-/-cells. **A**) CRISPR/Cas9 biallelic targeting strategy for *ORC5. ORC5* sgRNA target first methionine located upstream of Walker A motif. Second methionine locates at 133 aa. **B**) DNA sequences of wild type *ORC5* exon1 and three *ORC5Δ* clones obtained from genomic sequencing and by sequencing of cDNA. First methionine site is mutated in both allele of mutant clones. **C**) Verification of ORC5 antibody. Recombinant ORC5 protein with Myc tag were expressed and blotted with indicated antibodies. Ponceau S staining shows equal loading of lysate. **D**) Western blot of ORC5 in the *ORC5Δ* clones shows that full length and truncated ORC5 proteins are undetectable.

To ensure that an unexpected mRNA isoform of *ORC5* was not produced in these cells, the cDNA was amplified using primers matching exons 1 and 8 from *ORC5Δ* clones 19 and 24 and sequenced. The exon1 sequences of these two clones from genomic DNA or cDNA are identical and there was no evidence of alternative splicing between exons 1-8. No *ORC5* cDNA was amplified from clone101 suggesting that in this clone the mRNA is destabilized by the mutations in this clone.

### Three clones do not express any detectable ORC5 protein

ORC5 antibody was raised against full length of His tagged ORC5 protein. To test whether our ORC5 antibody detects short ORC5 protein translated from the 2^nd^ methionine at position 133, we expressed Myc tagged full length or short ORC5 recombinant protein (**Fig. 1C left**) and determined that both forms of ORC5 protein were detected by the ORC5 antibody (**Fig. 1C Middle**). Immunoblot with ORC5 antibody shows that clones 19, 24 and 101, do not express full length or N terminally deleted ORC5 (**Fig. 1D**). A quantitative immunoblot with dilutions of the cell lysates shows that the antibody can detect wild type level of ORC5 even in 6 μg of cell lysate, but cannot detect any protein in 120 μg of lysate from *ORC5Δ* cells (**Fig. 2A**). Thus if any ORC5 protein is produced in these clones, it is expressed at <5% the wild type level.

**Figure 2.**
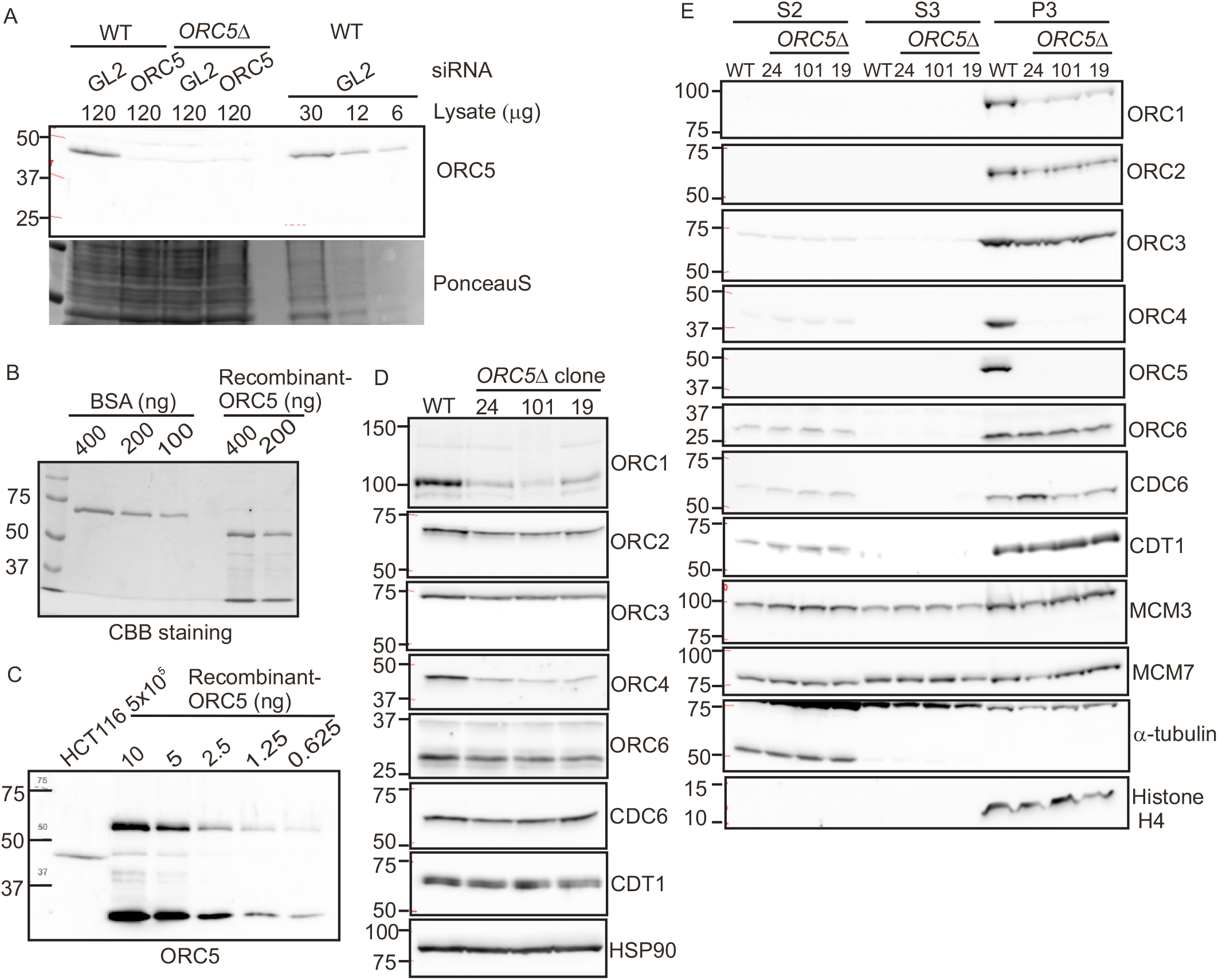
ORC protein in *ORC5Δ* cell lines. **A**) Quantitative Western blot for ORC5 shows that 6 μg of WT cell lysate contains enough ORC5 protein to be detected, yet 120 μg of lysate from *ORC5Δ* clones do not give a signal. So any undetectable ORC5 protein remaining in *ORC5Δ* cells is at <5% WT levels. Lane 2: Lysate from cells where ORC5 has been knocked down by siRNA shows the specificity of the anti-ORC5 antibody. GL2: negative control siRNA. **B**) Comparison of Coomassie Brilliant Blue staining of BSA and recombinant purified His-ORC5-AIM2PYD to show that the full length ORC5 protein (top band in recombinant ORC5 lane) is at 200 ng/μl. **C**) Immunoblot of HCT116 cell lysate with the top band of recombinant ORC5 to show that 5×10e5 WT cells give an ORC5 signal equal to 2.5 ng of recombinant ORC5, which corresponds to 240×10e8 molecules of ORC5. **D**) Immunoblot of indicated proteins in cell lysates of the WT and *ORC5Δ* clones. **E**) Immunoblot of soluble and chromatin-associated proteins in the WT and *ORC5Δ* clones.

To estimate how many molecules of ORC5 may remain undetected in the *ORC5Δ* cells, we titrated the antibody against purified recombinant His-ORC5-AIM2PYD. First, by comparing with BSA and Coomassie blue staining we determined how much volume of the recombinant protein contains 200 ng of His-ORC5-AIM2PYD (**Fig. 2B**). We then show that the immunoblot signal from 5×10^5^ HCT116 cells is similar to that from 2.5 ng of His-ORC5-AIM2PYD (**Fig. 2C**). Using a MW of 61.9 kD for His-ORC5-AIM2PYD and the Avogadro number, we can therefore calculate that ∼ 48,000 molecules of ORC5 are present per cell in WT HCT116 cells, so that our sensitivity of detection (<5% WT levels) is < 2400 molecules of ORC5/cell. Also note that even if a deleted ORC5 is expressed in the *ORC5Δ* cells at <2400 molecules/cell from an undetected splice isoform of the mRNA, it would have to use M133 for initiation and so the resulting molecule is expected to be nonfunctional.

### ORC4 and ORC5 are not loaded on the chromatin, but MCM2-7 is loaded on the chromatin in the *ORC5Δ* cells

The order of the ORC subunits in the ORC ring is: ORC2, ORC3, ORC5, ORC4, ORC1. ORC4 and ORC5 proteins are associated with each other in a bimolecular complex (22) and are adjacent to each other in the ORC ring (9). The loss of ORC5 is accompanied by the destabilization of ORC4 protein (**Fig. 2D**). Surprisingly, the ORC1 protein, not expected to be in contact with ORC5, is also destabilized, perhaps because of the decrease of ORC4. On the other hand, ORC3, which is in direct contact with ORC5, and ORC2 are not appreciably decreased in the cell lysates. As with the mutations in *ORC1* or *ORC2* (21), the mutation in ORC5 did not decrease the levels of ORC6, CDC6 or CDT1 in cell lysates (**Fig. 2D**) or the levels of MCM3 or MCM7 from the MCM2-7 ring in the soluble fractions of cells (**Fig. 2E**).

Cells were fractionated to determine the association of these proteins on chromatin. S2 and S3 are the soluble fractions while P3 is the chromatin associated fraction. The bulk of the ORC subunits even in WT cells is associated with the chromatin (**Fig. 2E**). In the *ORC5Δ* cells, ORC5 and ORC4 proteins were not detected on the chromatin fraction (**Fig. 2E**). However, ORC3 and ORC6 association with the chromatin was virtually unchanged, while ORC2 and ORC1 loading on the chromatin was moderately decreased, but still very detectable. Thus the six-subunit ORC does not have to form for some of the human subunits to be loaded on the chromatin, a finding consistent with our previous results (21), and at odds with the results in the yeasts where the six subunit holocomplex loads on the chromatin as one unit (23, 24). Even more surprising, given the role of ORC in recruiting CDC6 and CDT1+MCM2-7, we do not see any decrease of the chromatin association of CDC6, CDT1, MCM3 or MCM7 in the *ORC5Δ* cells.

### Complex formation between ORC subunits in the absence of ORC5

When ORC2 was deleted, all four subunits of the core ORC2-5 were destabilized making it difficult for us to test whether the remaining subunits still formed a core complex (21). In contrast, in the *ORC5Δ* cells, the persistence of ORC2, ORC3 and ORC6 allowed us to test whether at least two of the subunits of the core ORC subcomplex associated with each other in the absence of ORC5. Co-immunoprecipitation of ORC2 with ORC3 antibody in *ORC5Δ* cells suggest that at least those two ORC subunits can interact in the absence of ORC5 (**Fig. 3A**). Note that unlike in the yeasts, we and others have reported that ORC1 and ORC6 are very loosely associated with the ORC2-5 core even in WT human cells and so the association of ORC6 with ORC2+3 was not examined (25–27).

**Figure 3.**
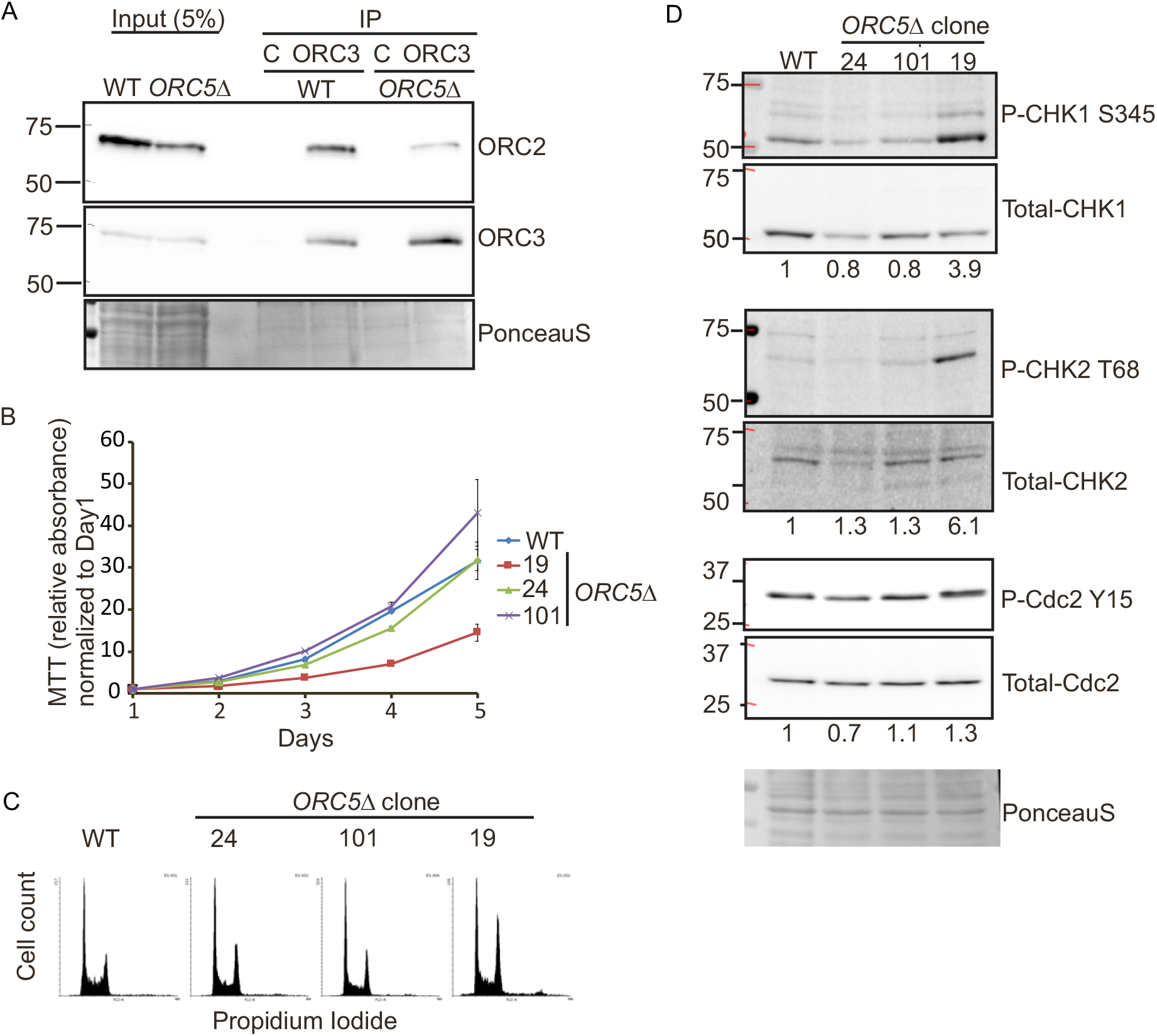
Cell proliferation in the *ORC5Δ* cell lines. **A**) Immunoprecipitation of ORC3 shows the co-precipitation of ORC2 in the *ORC5Δ* cells. C is control antibody. **B**) Cell growth of indicated clones over 5 days, expressed as MTT absorbance relative to the level at day 1. (Mean ± S.D.; n = 3 biological replicates). **C**) Cell cycle profile of propidium-iodide stained cell from indicated clones. **D**) Immunoblot of G2 checkpoint proteins in the WT and *ORC5Δ* clones. The numbers below each total protein were the ratio of phosphorylated protein and total protein.

### Cell proliferation and S phase in *ORC5Δ* cells

The proliferation rates of clones 101 and 24 were comparable to that of WT cells, while that of clone 19 was decreased, but still robust enough to expand the clone (**Fig. 3B**). The G2 fractions of *ORC5Δ* clone24 or 101 were comparable or slightly accumulated compared to WT cells in asynchronous cultures and *ORC5Δ* clone19 has apparent G2 accumulation (**Fig. 3C**). To test whether a defect in DNA replication triggers a G2 checkpoint in the *ORC5Δ* clones, we examined the phosphorylation of Chk1, Chk2, and Cdc2-Y15. Despite an increase of phosphorylation of Chk1 and Chk2 in *ORC5Δ* clone19, phosphorylation of Cdc2-Y15 is not as marked (**Fig. 3D**). Cdc2-Y15 phosphorylation is the end-result of the G2 checkpoint activation. The other two *ORC5Δ* clones do not show any sign of phosphorylation of Chk1, Chk2 or Cdc2-Y15. We conclude that the G2 checkpoint is not reproducibly activated in *ORC5Δ* cells. Consistent with this, when we followed the duration of S phase by releasing cells from an early S phase arrest with double-thymidine block, S phase was completed in *ORC5Δ* clone24 cells in the same 6-8 hr taken by WT cells (**Fig. 4**). The BrdU incorporation was diminished by a mere 25% in clones 24 and 101 (not tested for clone 19) (**Fig. 5A**). These results are comparable to what we observed in the cells without detectable ORC2 or ORC1 (21): cell proliferation and DNA replication are not as attenuated as would be expected if the six subunit ORC was absolutely essential for DNA replication.

**Figure 4.**
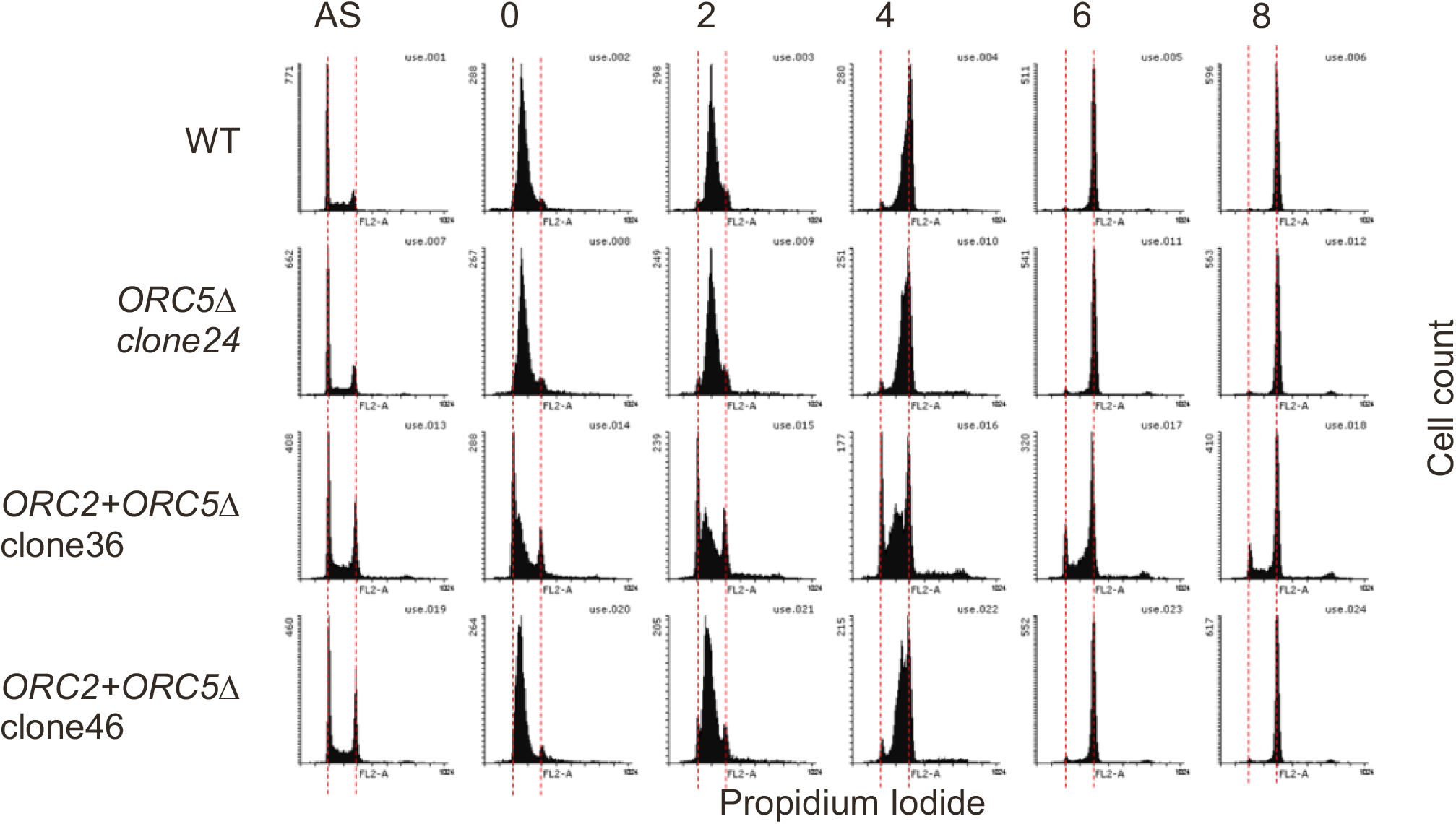
The duration of S phase in the *ORC5Δ* or *ORC2+ORC5Δ* clones. Early S phase arrested cells were released into nocodazole containing medium and collected at indicated time to measure the rate of S phase progression by propidium iodide FACS. AS: asynchronous cells. The red dotted lines indicate cell with 2N and 4N DNA content.

**Figure 5.**
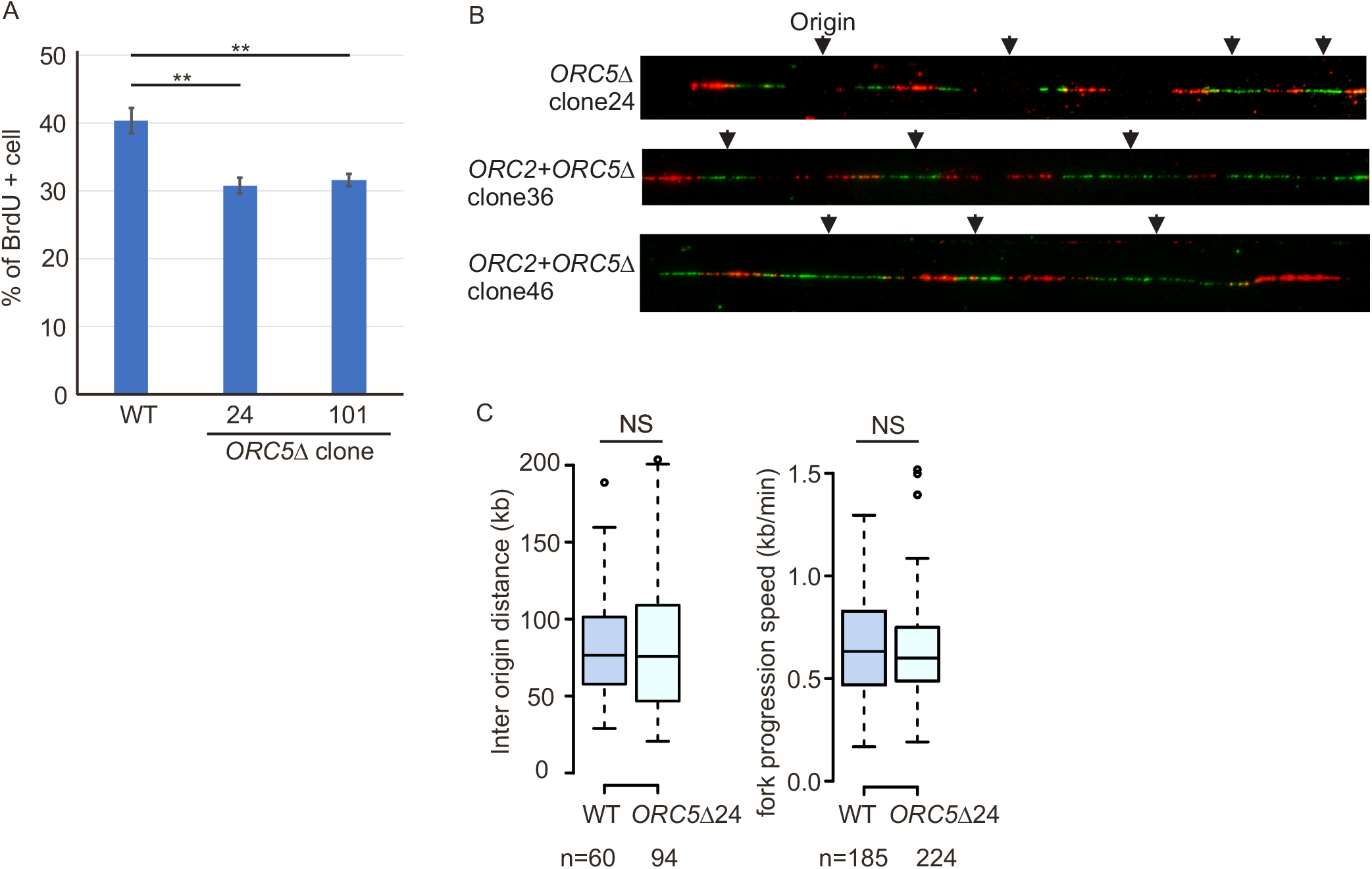
DNA replication in the *ORC5Δ* cell lines. **A**) Asynchronous cells were labeled with BrdU for 30 min. % of BrdU labeled cells were measured by two color FACS (P value < 0.01, two-sided t-test, Mean ± S.D. n = 3 biological replicates). **B**) Representative DNA combing images showing bi-directional origin firing in *ORC5Δ* or *ORC2*+*ORC5Δ* clones. DNA combing after a pulse labeling with IdU for 30 min followed by CldU for 30 min. **C**) Box and whisker plot for inter-origin distance (Left) and fork progression speed (Right) in DNA combing assays. NS: No significant difference between WT and *ORC5Δ* cells in two-sided Mann-Whitney U Test. N=number of tracks counted.

### Initiation of DNA replication in *ORC5Δ* cells

If fewer origins are licensed and fired in the ORC5 depleted cells, we expected to see an increase in inter-origin distance, and perhaps an acceleration of fork progression rates to compensate for the fewer active origins. However, molecular combing of DNA fibers obtained by labeling cells for 30 min with IdU followed by 30 min with CldU in *ORC5Δ* clone24 showed (a) that bidirectional origins were still being fired (**Fig. 5B**) and (b) there was no appreciable increase in the inter-origin distance in the cells with ORC5 depleted (**Fig. 5C**). There was also no increase in fork elongation rates. Thus there appears to be no decrease in the 52,000 origins that are fired per cell per S phase (21) even though the cells do not express functional ORC5 protein.

### Sensitivity of *ORC5Δ* cells to depletion of *CDC6*

To test whether the *ORC5Δ* cells are more dependent on CDC6, as we noted in *ORC1Δ* or *ORC2Δ* cells, we depleted CDC6 by siRNA and performed colony formation assays. Compared to WT cells, cell viability of *ORC5Δ* cells depleted of CDC6 is decreased, suggesting that CDC6 function becomes more important when the ORC5 subunit is deleted (**Fig. 6**).

**Figure 6.**
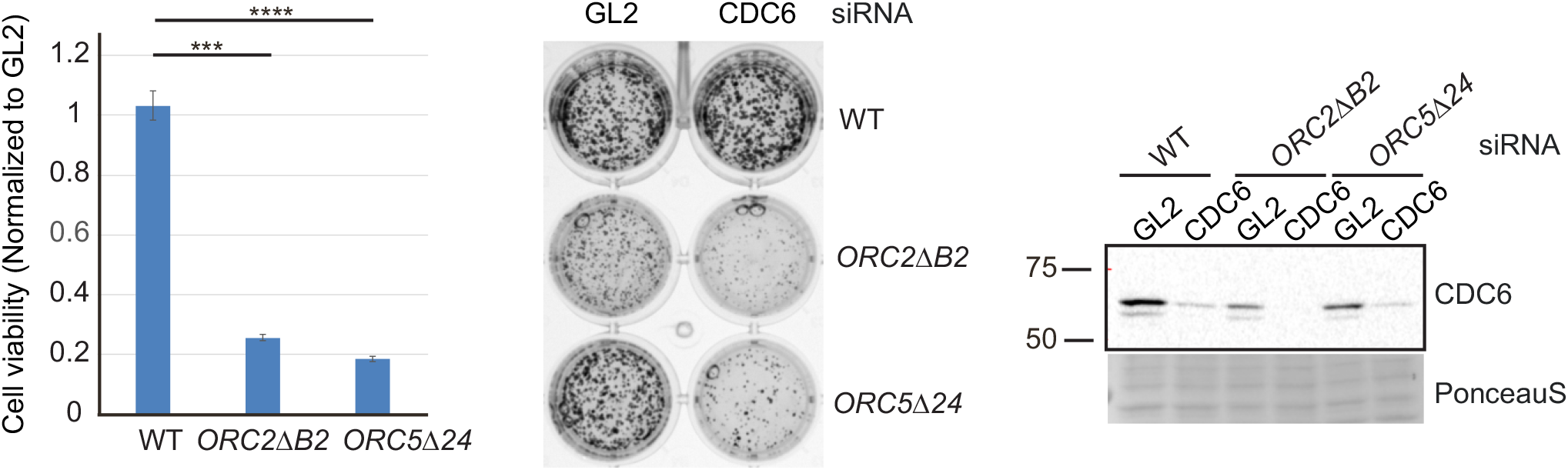
*CDC6* is more essential for cell viability in the *ORC5Δ* cells. Right: Western blot of lysate from indicated cells transfected with siGL2 (negative control) or siCDC6. Middle: 24 hours after transfection, 1000 cells were re-plated for Colony formation and stained by CellTiter 96 after 5 days. Left: Colony density were measured. Data presented for each cell line normalized to the density of the siGL2 transfected cells. (****P value < 0.00001, *** P value <0.0001, two-sided t-test, Mean ± S.D. n = 3 biological replicates).

### Biallelic disruption of the *ORC5* and *ORC2* genes

So far, we showed that cells where a single ORC subunit (ORC1, ORC2 or ORC5) is depleted by mutation can proliferate and replicate ((21)and here). To test whether more than two ORC subunits can be deleted in a cancer cell line, we deleted *ORC2* simultaneously in the *ORC5Δ* cells. Since ORC3 or ORC4 proteins require ORC2 or ORC5 protein, respectively, for their stability and integration into the ORC complex we expected that several subunits of the ORC ring will become undetectable in these cells. We used CRISPR/Cas9 to introduce frame shift mutations in both alleles of exon 4 of *ORC2* gene in *ORC5Δ* clone24. After ORC2 sgRNA transfection into *ORC5Δ* cells, cells were plated in 96 well plates and screened by western blot to identify clones where ORC2 and ORC5 proteins are undetectable. We confirmed that the *ORC2+ORC5Δ* clones express neither ORC2 nor ORC5 protein by western blotting (**Fig. 7A**). No band was detected even by an antibody recognizing the C-terminal part of ORC2 suggesting that even a short form of ORC2 protein is not expressed in these clones (**Fig. 7A**). In our previous work we showed that the sensitivity of detection of ORC2 protein was at ∼1500 molecules/cell, so that if any ORC2 protein was expressed at levels below our detection threshold, the level is still <1500 molecules/cell. In addition to western blotting, we also validated the biallelic mutation of *ORC2* by sequencing the genomic DNA and the cDNA of *ORC2* (**Fig. 7B**).

**Figure 7.**
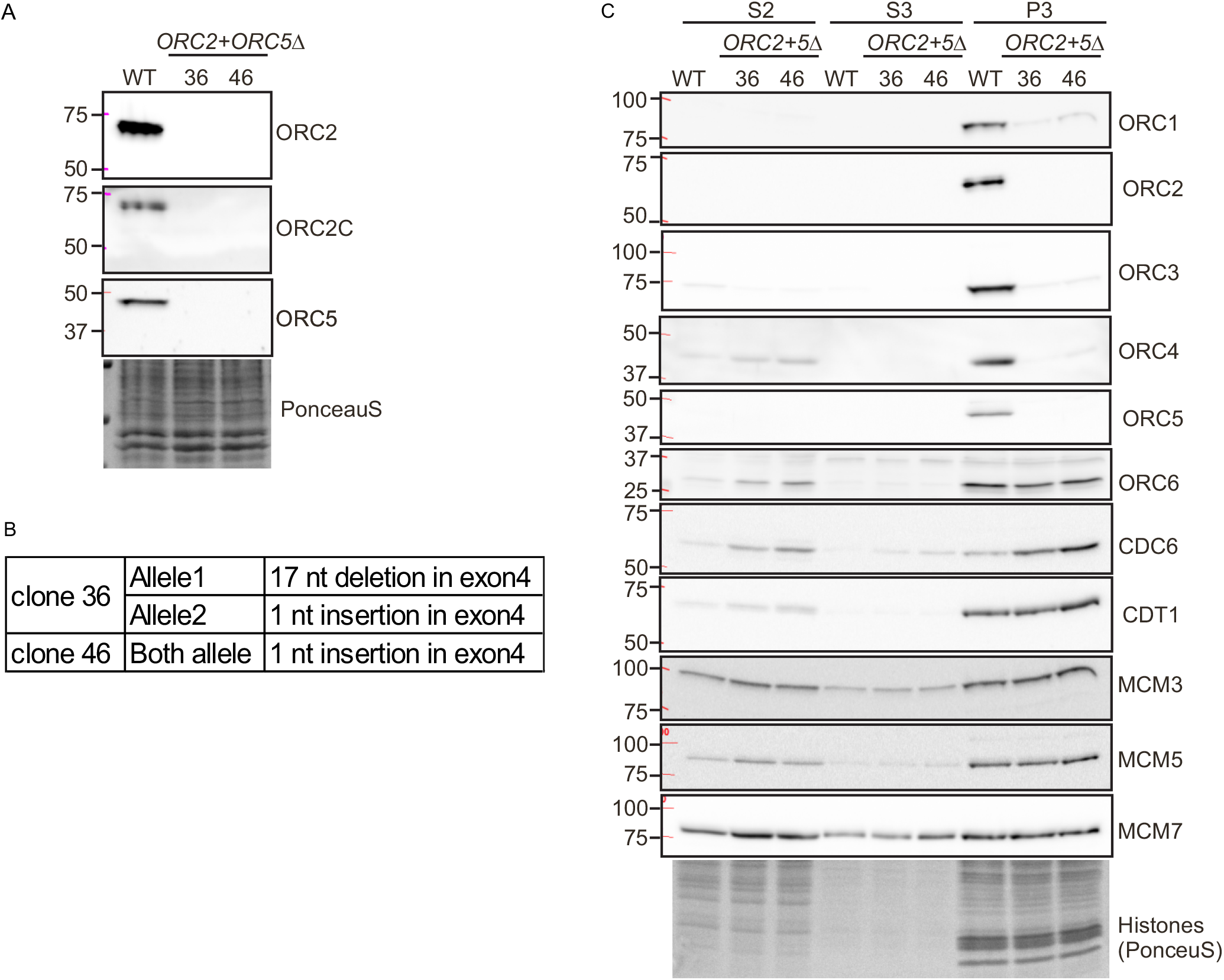
Simultaneous deletion of *ORC2* and *ORC5* in HCT116 p53-/-cells. **A**) Western blot of ORC2 and ORC5 in the *ORC2+ORC5Δ* clones. **B**) DNA sequences of *ORC2* exon 4 show biallelic frame-shift mutation of *ORC2* in the *ORC2+ORC5Δ* clones. **C**) Immunoblot of soluble and chromatin-associated proteins in the *ORC2+ORC5Δ* clones.

### Unchanged chromatin loading of MCM2-7

Again, subcellular fractionation of WT cells show that the bulk of the ORC subunits was associated with the chromatin fraction **(Fig. 7C)**. In addition to the ORC2, 3, 4 and 5 subunits, ORC1 was also undetectable on the chromatin in *ORC2*+*ORC5Δ* clones (**Fig. 7C**). Similar to what was seen in the ORC5 depleted cells, MCM3, MCM5 and MCM7 loading on chromatin was unchanged in *ORC2*+*ORC5Δ* clones. This result was surprising because deletion of *ORC2* alone significantly reduced the chromatin loading of MCM2-7 complex (21). The *ORC5Δ* clones must have acquired some mechanism to efficiently recruit MCM2-7, and this mechanism persists in the *ORC2*+*ORC5Δ* clones (**Fig. 7C**). As before with the clones mutated individually in *ORC1, ORC2* or *ORC5*, ORC6 and CDT1 association with chromatin was unchanged in the double deletion cells. CDC6 association with chromatin was significantly increased in the double deletion cells (1:4:10 for WT: clone 36: clone 46). Combined with previous results showing that ORC subunit-deleted cells are more sensitive to CDC6 depletion ((21) and **Fig. 6**), the increase in CDC6 association with chromatin in the *ORC2Δ* (21) and *ORC2+ORC5Δ* cells suggests that CDC6 may compensate for the function of the impaired ORC complex in loading MCM2-7.

### Cell proliferation and DNA replication in *ORC2*+*ORC5Δ* cells

*ORC2*+*ORC5Δ* cells grew slower than WT or parental *ORC5Δ* clone 24. However the cells still proliferate and are viable after over 2 months. (**Fig. 3B, 8A**), and cell cycle profiling by FACS analysis shows that both *ORC2*+*ORC5Δ* clones show accumulation of cells in G2/M phase (**Fig. 8B**). To test whether the additional mutation of *ORC2* in *ORC5Δ* cells slows the progression of S phase, cells were synchronized in early S phase by double thymidine block and released into nocodazole containing medium to arrest cell in M phase (**Fig. 4**). WT, parental *ORC5Δ*, and *ORC2*+*ORC5Δ* clone 46 show a similar duration of S phase, suggesting that the slow cell proliferation in that *ORC2*+*ORC5Δ* clone is not due to the impairment of S phase progression. *ORC2*+*ORC5Δ* clone 36 may have a slight prolongation of S phase, but as can be seen in **Fig. 8A**, this clone actually shows a slightly faster proliferation than clone 46, and so we conclude that the slow cell proliferation in either of the double deletion clones is not due to the prolongation of S phase

Finally, we performed a DNA combing assay to characterize the DNA replication initiation and fork elongation in the *ORC2*+*ORC5Δ*. Despite undetectable chromatin loading of ORC1-5 subunits, bidirectional origins of replication were still detected (**Fig. 5B**) and the *ORC2*+*ORC5Δ* clone36 licensed and fired more origins (with shorter inter-origin distance) and fork progression rate was slower than WT cells (**Fig. 8C**). The second clone has a longer inter-origin distance and faster fork progression (**Fig. 8C**). Because the two clones have opposite effects on inter-origin distance, we conclude that despite the double deletion these cancer cells can fire near wild type levels of origins (52,000 per cell per S phase). Thus origin licensing and initiation can carry on at near normal rates even if ORC2 and ORC5 proteins are completely undetectable and three of the other subunits of the ORC ring are also not detectable on chromatin.

**Figure 8.**
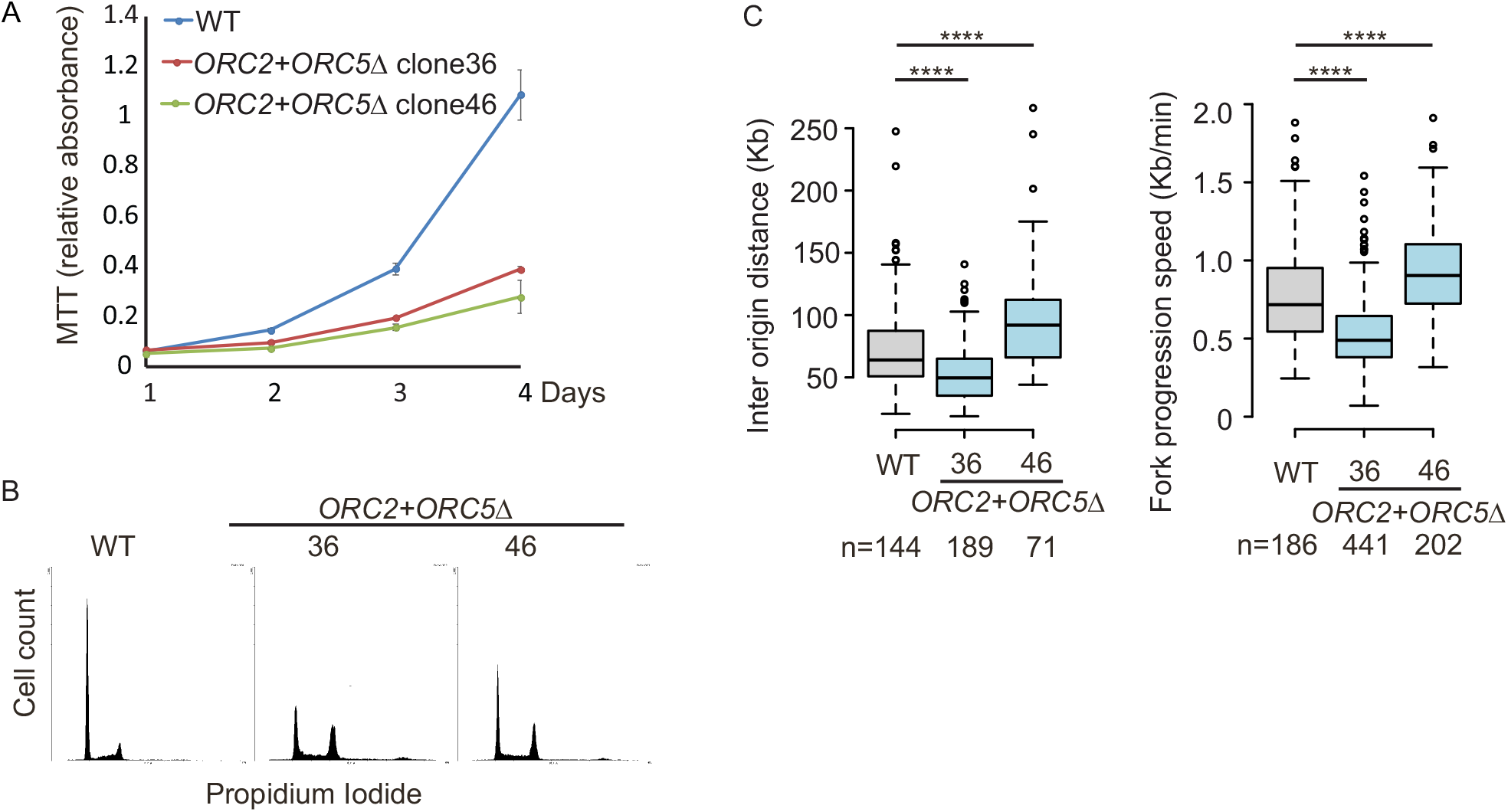
Cell proliferation and DNA replication in the *ORC2*+*ORC5Δ* cell lines. **A**) Cell growth of indicated clones over 4 days, expressed as MTT absorbance. (Mean ± S.D.; n = 3 biological replicates). **B**) Cell cycle profile of propidium-iodide stained cells from indicated clones. **C**) DNA combing after a pulse labeling with IdU for 30 min followed by CldU for 30 min. Box and whisker plot for inter-origin distance (Left) and fork progression speed (Right) (****P<0.00001 between WT and *ORC2*+*ORC5Δ* clones in two-sided Mann-Whitney U Test; N=number of tracks counted.).

## Discussion

When we reported that cancer cell lines could proliferate and replicate in the absence of detectable levels of two subunits of ORC, ORC1 and ORC2, we suggested that “the absolute requirement for six-subunit ORC for licensing bi-directional origins of replication can be bypassed in some cell lines”. We were careful to point out that this does not meant that all ORC subunits are dispensable, or that ORC will not be essential in most circumstances for initiation of DNA replication. Our results only showed that it was possible to select for cancer cell lines that could replicate in the apparent absence of ORC1 or ORC2, though there was always the caveat that splice isoforms, or an internal translation initiation site produces small amounts of truncated ORC1 or ORC2 protein that was below the level of detection, but sufficient for origin licensing.

In the present work, we confirmed by cloning and sequencing the cDNA of *ORC5*, that the mutant *ORC5* allele could not produce an ORC5 protein with an intact AAA+ ATPase domain, even at levels below detection. Any ORC5 protein produced, would have had to initiate from a methionine that is downstream from the Walker A motif, which would have deleted most of the ATPase lobe of ORC5. Even if initiation began independent of a methionine, the quantitative immunoblot results show that not much ORC5 protein is being generated (<2400 molecules/cell). Yet, the cells recruit MCM2-7, initiate approximately the same number of bidirectional origins as wild type cells (∼52,000 per cell per cell-cycle (21)) and replicate their chromosomes and proliferate fairly normally.

In many respects the phenotype of the cells surviving without ORC5 protein are milder than the phenotypes of the cells surviving without detectable ORC2 protein (17). ORC5 loss primarily destabilizes ORC1 and ORC4 alone, while ORC2 loss de-stabilized ORC1, 3, 4 and 5 and CDT1. In both mutants, ORC6, CDC6 and CDT1 recruitment to chromatin was relatively unimpaired, and MCM2-7 was loaded sufficiently for normal DNA replication. These observations contradict the original model derived from studies in yeast and in Xenopus egg extracts that a six–subunit ORC has to be present on DNA to facilitate the loading of CDC6, CDT1 and MCM2-7. Instead it appears that some subunits of ORC, e.g. ORC6, associate with chromatin independent of the other subunits, and that CDC6 or CDT1 can be recruited to chromatin independent of many of the ORC subunits. It is also possible that the bulk of the chromatin binding seen for ORC6, CDC6 or CDT1 is not due to their functions at origins of replication, but due to as yet undiscovered functions independent of origin licensing. Although DNA replication is near normal in the *ORC2Δ* cells, MCM2-7 binding to chromatin was more sensitive to ORC2 loss in the *ORC2Δ* cells than ORC5 loss in the *ORC5Δ* cells. One possible bypass mechanism in the cells mutant in individual ORC subunits is that a crippled ORC lacking ORC5 and ORC4 (in the *ORC5Δ* cells) or ORC2 and ORC3 (in the *ORC2Δ* cells) can recruit MCM2-7. However, the results with the *ORC2+ORC5Δ* clones reported here, where five of the six subunits of ORC are undetected on chromatin, and yet MCM2-7 binding is unaffected and nearly wild type levels of bidirectional origins of replication fire, make it more likely that MCM2-7 recruitment does not always need the ORC1-5 ring or even a partial ORC ring.

By itself, the *ORC2Δ* (21) decreased MCM2-7 loading on chromatin, but not sufficiently to impair origin firing. However, here when *ORC2* is deleted in cells that have already adjusted to the loss of ORC5, MCM2-7 association with chromatin is virtually at wild type levels. This could be explained if ORC5 acts as a repressor of MCM2-7 recruitment, so that its presence in *ORC2Δ* cells is deleterious for MCM2-7 binding to chromatin. The other possibility is that the *ORC5* deletion allowed cells to acquire a robust pathway to bypass ORC for MCM2-7 recruitment, a pathway that was not activated when *ORC2* was deleted by itself.

ORC2-3-5-4-1 form a gapped ring (with a central channel that is wide enough to surround a DNA double-helix) with the subunits arranged around the ring in the order specified. Later, during the loading of MCM2-7 ring, CDC6 is hypothesized to fill the gap between ORC2 and ORC1. Modeling on the interaction of the RF-C clamp loader on the PCNA ring, the ORC-CDC6 ring is believed to interact with the MCM2-7 ring end-on-end to facilitate the opening of the MCM2-7 ring so that it can be loaded around the DNA (9, 10, 23, 28, 29). We suspect that after loss of two subunits (ORC2 and ORC5) in the same cell, the remaining ORC subunits will be unable to make a ring large enough (i) to encircle a DNA double-helix or (ii) to interact with the MCM2-7 ring end-on-end. Of course this problem can be circumvented if some subunits are used multiple times in the ring. But even with that strategy, the destabilization of three of the remaining subunits seen in the double-mutant cells leaves very few subunit molecules to form enough ORC rings to load near normal levels of MCM2-7 on chromatin by clamp-loading activity. ORC6, of course, is very different in sequence from the other five subunits and has not been located in the ORC2-3-5-4-1 ring, making it unlikely to form a homomeric ring. The other alternative is that some isolated ORC subunits, perhaps ORC6 alone, helps recruit MCM2-7 to the chromatin, but the ORC ring is not necessary to open the MCM2-7 ring and load it around the DNA.

We next turn to whether human MCM2-7 can be recruited to chromatin without the ATPase activity of ORC. Unlike the situation in the yeasts where ORC1-5 subunits have Walker A and B motifs, ORC1 and ORC4 are the only subunits of human ORC with intact Walker A and B motifs. Furthermore, the Walker A and B motifs of the ORC1 subunit seem to be the only motifs essential for the ATPase activity of ORC in *S. cerevisiae, D. melanogaster* and *H. sapiens* and this ATP binding and hydrolysis activity is essential for ORC function (18–20). In our previous paper we found the cells survive with mutations that make ORC1 protein undetectable (21). Now we find in the *ORC2*+*ORC5Δ* cells that ORC1 and ORC4 proteins are undetectable on chromatin, and yet MCM2-7 is bound at near wild-type levels to chromatin, raising the possibility that MCM2-7 can be recruited to chromatin to form a functional replicative helicase without the ATPase activity of ORC. This is consistent with recent reports that the ATPase activity of yeast ORC is not required for MCM2-7 loading (13).

The clamp-loader model of ORC+CDC6 in loading MCM2-7 is modeled after the similarities of the RF-C ATPase ring serving as a loader of the ring-shaped PCNA clamp. The suggestion is that the ATPase activity of the ORC+CDC6 loader cracks open the MCM2-7 ring to enable it to encircle the DNA. Our results suggest that this may not be absolutely essential for human replication initiation. One possibility is that the MCM ring is kept in a cracked state by CDT1 (14, 15) or is already a pre-cracked ring as in archaea where it is recruited to DNA by a single protein that is related to both Orc1 and Cdc6 (16). Therefore, the primary function of ORC may be to recruit MCM2-7 to sites near the DNA, and that MCM2-7 can encircle the DNA without the benefit of a clamp-loader. Of course, our results also suggest that even the recruitment of MCM2-7 to chromatin is not absolutely dependent on the six-subunit human ORC. Here some combination of ORC6, CDC6 and CDT1 may be sufficient to recruit MCM2-7 to the DNA in the *ORC5Δ* or *ORC2+ORC5Δ* cells at a level that is comparable to that in WT cells.

There are a few examples in the Literature that have also suggested the possibility that in certain circumstances the replicative helicase in higher eukaryotes may be loaded on DNA in the absence of the six-subunit ORC. *ORC1-/-* Drosophila larvae still allowed endoreplication in salivary gland cells, with only a twofold reduction of ploidy (30). Conditional deletion of mouse *ORC1* shows that the protein is essential for viability of embryonic cells and for proliferation of intestinal epithelial cells (7). However, the mice continued to endoreplicate the nuclei in the liver during liver regeneration, again suggesting that occasionally mammalian cells can load enough MCM2-7 on chromatin for successful chromosomal replication in the absence of the six subunit ORC.

Given that MCM2-7 recruitment to chromatin and origin firing appears near normal in the *ORC2+ORC5Δ* cells, what accounts for the decrease in cell proliferation and the accumulation of cells in G2/M? Subunits of ORC have also been implicated in centrosome biology or cytokinesis (31–33) so it would be interesting to examine whether these functions account for slow proliferation. In addition, we are currently investigating whether other functions of ORC, like its role in epigenetics (34–36), is responsible for the decrease in proliferation despite the near normal replication initiation. As we have noted, decrease of ORC subunits decreases cell proliferation without affecting replication initiation. This observation should be borne in mind when interpreting standard siRNA or sgRNA screens for viability of cells after ORC depletion. Such screens may indicate that ORC subunits are important for cell proliferation, but unless tested specifically by molecular combing assays for an increase in inter-origin distance, cannot say whether replication initiation is impaired.

In summary, we suggest that the absolute requirement for a six-subunit ORC for licensing bi-directional origins of replication can be bypassed in some circumstances. The increased dependence of the *ORC* mutant cells on CDC6 for viability makes it likely that CDC6, perhaps with ORC6, can under exceptional circumstances carry out the function of the ORC ring in recruitment and loading of the MCM2-7 pre-helicase complex around DNA.

## Experimental procedures

### Cell culture and transfection

HCT116 p53−/− cells (37)were maintained in McCoy’s 5A-modified medium (Corning Inc., Corning, NY) supplemented with 10 % (v/v) fetal bovine serum (Thermo Fisher Scientific) and penicillin/streptomycin. The plasmids were transfected with Lipofectamine 2000 (Thermo Fisher Scientific, Waltham, MA) and siRNAs transfected with RNAiMAX (Thermo Fisher Scientific) according to the manufacturers” protocol. CDC6 siRNA (GAUCGACUUAAUCAGGUAU) was synthesized by Thermo Fisher Scientific. To measure the duration of S phase progression, cells were synchronized at early S phase by Double thymidine block. In brief, 2 mM thymidine was added for 18 hr. and removed for 8 hr. before adding 2^nd^ round of 2 mM thymidine for 18hr. Early Synchronized cells were washed and released into fresh media containing 100ng/ml nocodazole to trap in M phase and collected at various time point. No mycoplasma contamination was found. HCT116 p53-/-cells were authenticated by STR profiling (Biosynthesis, Lewisville, TX).

### Plasmids

ORC5 sgRNA (GTTTTCCAAGTGGGGCATTC) was cloned into p413-Cas9 vector backbone (a generous gift from Adli Lab, Northwestern University) using PCR and In-Fusion cloning (Clontech, Mountain View, CA). ORC2 sgRNA (GAAGGAGCGAGCGCAGCTTT) was cloned into pCR-Blunt II-TOPO vector backbone (Addgene 41820, Cambridge, MA) using PCR and In-Fusion cloning (Clontech). Human codon optimized Cas9 nuclease (hCas9) was used (Addgene (41815)). To express Myc tagged ORC5 in mammalian cells, *ORC5* was cloned into pEFM vector that is under EF1α promoter and expresses a N-terminal Myc tag using PCR and In-Fusion.

### DNA combing assay

Cells were pulse-labeled for 30 min each and in succession, with 100 μM 5- chlorodeoxyuridine (ldU), followed by 250 μM 5-iododeoxyuridine (CIdU) and embedded into agarose plugs. DNA was combed on silanized coverslips (Genomic Vision, Bagneux, France), dehydrated at 65 °C for 4 hr., denatured in 0.5 M NaOH and NaCl for 8 min, and dehydrated in a series of ethanol concentrations. CldU or IdU were detected by immunofluorescence with either anti-BrdU antibody that recognizes CldU (MCA2060, Bio-Rad Laboratories, Hercules, CA) or anti-BrdU antibody that recognizes IdU (347583, BD Biosciences, San Jose, CA). Images were acquired on a Zeiss AxioObserver Z1, 63 X objective and DNA lengths measured using ZEN software.

### BrdU incorporation

Bromodeoxyuridine (BrdU) incorporation was measured as previously described (38) with minor modifications. Cells were labeled with 10 μM BrdU for 30 min and fixed in 70 % Ethanol. DNA was denatured in 2 M hydrochloric acid and stained with FITC-conjugated BrdU antibody (556028, BD Biosciences) and propidium iodide (MilliporeSigma, St. Louis, MO) according to the manufacturer’s instruction.

### Clonogenic assay

Cells were transfected with siRNA to CDC6. After 24 hours, 1000 cells were plated in 24 well plates. 5 days later, Colonies were stained with CellTiter 96® Non-Radioactive Cell Proliferation Assay (Promega, Fitchburg, WI) and recorded with plate reader. All experiments were conducted in triplicate.

### Proliferation assay

1000 cells were plated per well in 96 well plates. The absorbance of cells was measured every 24 hours using CellTiter 96® Non-Radioactive Cell Proliferation Assay (Promega, Madison, WI) according to the manufacturer’s instructions. All experiments were conducted in triplicate.

### Immunoprecipitation, western blot, and antibodies

Cells were lysed in lysis buffer 50 mM Tris-HCl (pH 8.0), 150 mM NaCl, 5 mM EDTA, 0.5 % NP-40, 1 mM DTT, 20 mM NaF, and protease inhibitor cocktail. Lysate was cleared by centrifugation and incubated with anti-ORC3 antibody (sc-23888, Santa Cruz Biotechnology, Dallas, TX) for 4 hours. Immunoprecipitate was collected on Dynabeads Protein G (Thermo Fisher Scientific), washed in lysis buffer and eluted with 2 x SDS sample buffer. Antibodies used were as follows. ORC1 (4731, Cell Signaling Technology, Danvers, MA), ORC2 (sc-32734, Santa Cruz Biotechnology), ORC2C (sc-13238, Santa Cruz Biotechnology), ORC3, ORC4, ORC5, and ORC6 (38), MCM3 (sc-9850, Santa Cruz Biotechnology), MCM5 (A300-195A, Bethyl Laboratories, Montgomery, TX), MCM7 (sc-9966, Santa Cruz Biotechnology), CDC6 (3387, Cell Signaling Technology), Phospho- Chk1 (Ser345, 2348, Cell Signaling Technology), Phospho-Chk2 (Thr68, 2661, Cell Signaling Technology), Phospho-cdc2 (Tyr15, 9111, Cell Signaling Technology), Chk1 (NB-100-464, Novus Biologicals), Chk2 (sc-5278, Santa Cruz Biotechnology), Cdc2 (9116, Cell Signaling Technology), and Cdt1 (39), alpha-tubulin (sc-5286, Santa Cruz Biotechnology), Histone H4 (07-108, MilliporeSigma), HSP90 (sc-13119, Santa Cruz Biotechnology), MYC (9E10, homemade).

### Recombinant proteins

*ORC5* cDNA was cloned into pET28 AIM2PYD backbone (40) (a generous gift from Hao Wu, Harvard) with PCR and infusion cloning. His-ORC5-AIM2PYD (61.9 kD) protein with His-tag located at the N-terminus was expressed in *E*.*Coli* BL21 in LB media. Culture was induced with 0.1 mM of IPTG for 2 hours at 30 °C. For purification, pellet was resuspended in Guanidine buffer (100 mM sodium phosphate, [pH8.0], 10 mM Tris-HCl [pH8.0], and 6 M GuHCl). Lysate was incubated with Ni-NTA agarose (Qiagen, Hilden, Germany) and wash with Urea buffer (100 mM sodium phosphate [pH8.0], 10 mM Tris-HCl [pH8.0], 8 M UREA, and 10 mM Imidazole) before eluting with 0.4 M imidazole.

### Chromatin fractionation

Chromatin fractionation was performed as previously described (41). Cells were resuspended in buffer A (10 mM HEPES [pH7.9], 10 % glycerol, 1 mM DTT, protease inhibitor cocktail [Thermo Fisher Scientific]). After adding 0.1% Triton X-100, cells were incubated for 5 min on ice and centrifuge at 1300g, 4°C. Supernatants were clarified by centrifugation at 20000g, 4°C (S2). Pellets (Nuclei) were washed in buffer A and lysed in buffer B (3 mM EDTA, 0.2 mM, EGTA, 1 mM DTT, protease inhibitor cocktail). After 30 min on ice, lysate was centrifuged at 1700g, 4°C and supernatants were collected (S3). Pellets (chromatin) were washed in buffer B and lysed in 2 x SDS sample buffer and sonicated (P3).

## Statistical Analysis

Mann-Whitney U Test or two-sided t-test was performed to test the difference.

## Acknowledgments

We thank Dutta lab members for the useful discussion.

## Author Contributions

A.D. and E.S. initiated and designed the study. E.S. performed the experiments. The manuscript was prepared by A.D. and E.S.

## Funding and additional information

This study is supported by R01 CA060499 (to A.D.). The content is solely the responsibility of the authors and does not necessarily represent the official views of the National Institutes of Health.

